# Wavelength-dependent DNA photodamage in a 3-D human skin model over the far-UVC and germicidal-UVC wavelength ranges from 215 to 255 nm

**DOI:** 10.1101/2021.12.14.472653

**Authors:** David Welch, Marilena Aquino de Muro, Manuela Buonanno, David J Brenner

## Abstract

The effectiveness of UVC to reduce airborne-mediated disease transmission is well-established. However conventional germicidal UVC (~254 nm) cannot be used directly in occupied spaces because of the potential for damage to the skin and eye. A recently studied alternative with the potential to be used directly in occupied spaces is far-UVC (200 to 235 nm, typically 222 nm), as it cannot penetrate to the key living cells in the epidermis. Optimal far-UVC use is hampered by limited knowledge of the precise wavelength dependence of UVC-induced DNA damage, and thus we have used a monochromatic UVC exposure system to assess wavelength-dependent DNA damage in a realistic 3-D human skin model. We exposed a 3-D human skin model to monowavelength UVC exposures of 100 mJ/cm^2^, at UVC wavelengths from 215 to 255 nm (5-nm steps). At each wavelength we measured yields of DNA-damaged keratinocytes, and their distribution within the layers of the epidermis. No increase in DNA damage was observed in the epidermis at wavelengths from 215 to 235 nm, but at higher wavelengths (240-255 nm) significant levels of DNA damage were observed. These results support use of far-UVC light to safely reduce the risk of airborne disease transmission in occupied locations.

## INTRODUCTION

Ultraviolet (UV) radiation encompasses wavelengths from 100 nm to 400 nm, and is further categorized into UVC (100-280 nm), UVB (280-315 nm), and UVA (315-400 nm). The effectiveness of UVC radiation to inactivate or kill microbes in the air, on surfaces, or within liquids is well-established (1). Epidemiological studies by Wells *et al*. in the 1930s and 1940s demonstrated the ability of UVC installations to effectively reduce the transmission of airborne diseases (2), and upper-room ultraviolet germicidal irradiation remains an effective technology which is in use internationally (3).

However, use of conventional germicidal UVC (254 nm) fixtures is limited to exposing unoccupied spaces, such as the upper-room air volume, because of the potential health hazards associated with direct exposure to this wavelength to the skin or eye, respectively through erythema or photokeratitis (4, 5).

A recent alternative to 254 nm conventional germicidal UVC is far-UVC (wavelength range from 200 to 235 nm, typically used at 222 nm). Far-UVC is designed to be used directly in occupied indoor locations, with good evidence published both for efficacy to inactivate airborne pathogens including influenza and coronavirus (6–15), and safety for human exposure (16–20). Far-UVC safety is premised on the fact that, because its effective range in biological material is much shorter than for conventional (254 nm wavelength) germicidal UVC (16, 21–23), far-UVC incident on the skin is absorbed primarily in the superficial stratum corneum (see Fig. 1, containing only dead cells) and, to a much lesser extent in the adjacent stratum granulosum (granular layer, see Fig. 1, containing dead or dying cells moving to the stratum corneum). Far-UVC light is not expected (16, 21) to penetrate to the deeper stratum spinosum (spinous layer, see Fig. 1) or to the still deeper stratum basale (basal cell layer, see Fig. 1) of the epidermis, where DNA damage can result in long-term sequelae including carcinogenesis (24, 25). Similar considerations apply for the eye with regard to the tear layer and the superficial cells of the cornea. In term of efficacy, however, because of the small size of viral and bacterial pathogens, far-UVC can penetrate and inactivate these pathogens, typically with similar or improved efficacy compared with conventional (254 nm) germicidal UVC light (26).

**Figure 1.**
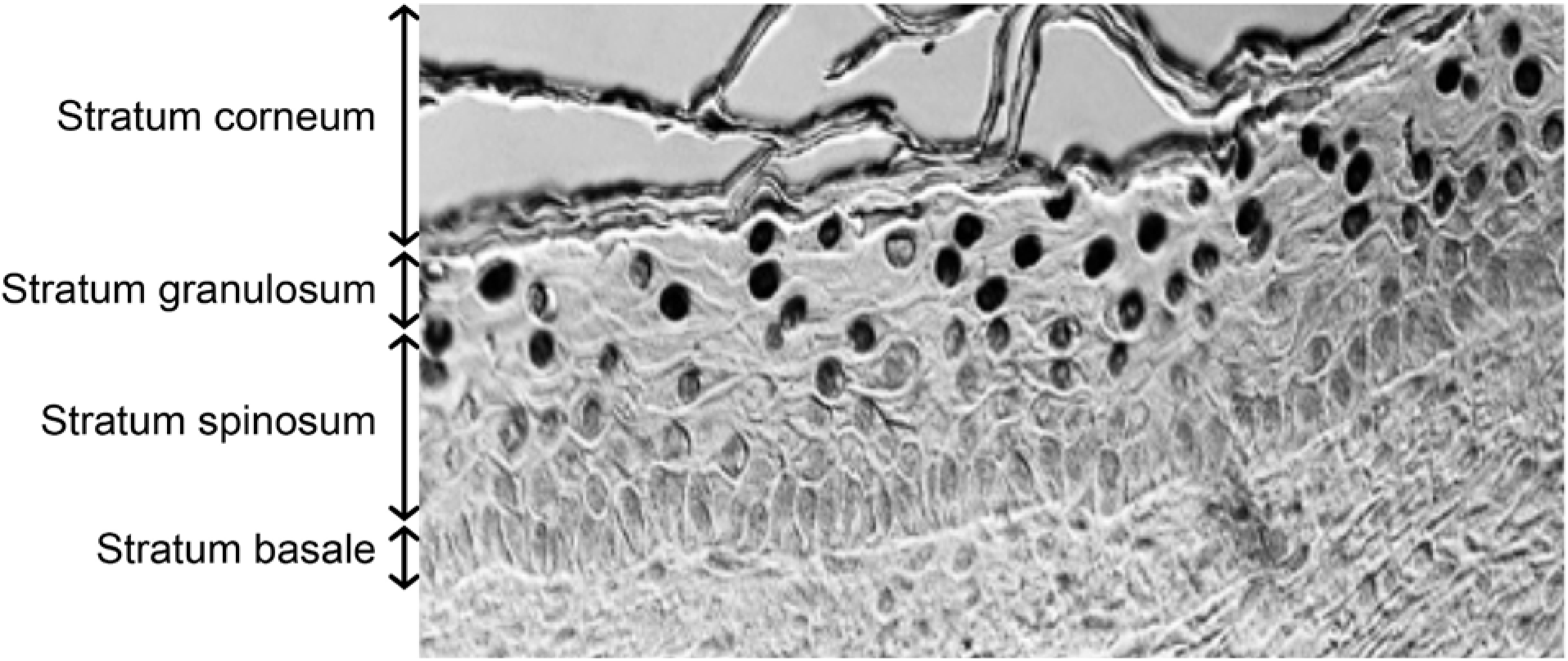
Representative image of the different layers of the epidermis in the 3-D human skin model, in this case exposed to 250 nm wavelength UVC. Cells with CPD DNA photodamage appear as dark-stained nuclei,

While there is considerable evidence for far-UVC safety in skin and eyes (7, 16, 18–20, 22, 27–31), there have been no direct systematic measurements of DNA damage in skin as a function of wavelength that encompasses the far-UVC and conventional germicidal UVC wavelengths. This is important both from the perspective of directly validating the far-UVC concept, but also because in addition to the primary emission (for example from a KrCl* excimer lamp at 222 nm) all far-UVC light sources also emit small fluences of higher wavelength UVC. These associated higher wavelength UVC emissions have been shown to result in DNA damage (17), and thus most far-UVC light sources use filters to remove them. Understanding the wavelength dependence of DNA damage will allow more efficient safe filters to be designed.

Our final rationale for this study is to contribute towards improved recommendations of the UVC action spectrum and associated exposure limits, which are currently under review [4, 39] by the ACGIH (American Conference of Governmental Industrial Hygienists) and the ICNIRP (International Commission on Non-Ionizing Radiation Protection), the agencies which provide regulatory recommendations in regard to UV Threshold Limit Values or Exposure Limits.

In this study, we used a monochromatic exposure system designed for narrow bandwidth UVC exposures, with which we irradiated realistic 3-D models of human skin which recapitulates the key components of human skin. Using this system we assessed the wavelength dependence of DNA photodamage measured in the whole epidermis and within the different epidermal layers.

## MATERIAL AND METHODS

### Monochromatic wavelength UVC exposure system

An optical system was assembled to enable monochromatic UVC exposures to 3-D models of human skin tissue. An EQ-77 Laser-Driven Light Source (Energetiq Technology, Inc., Wilmington, MA) provided a high brightness broadband output across the wavelength range of 170 nm – 2500 nm. A pair of off-axis parabolic mirrors focused the EQ-77 output into a Cornerstone 260 1/4 m monochromator (CS260-RG-2-FH-A, Newport, Irvine, CA). The monochromator was equipped with a 1201.6 g/mm plane blazed holographic reflection grating (#200H with master no. 5482, Newport) to maximize optical throughput in the UVC. Fixed slits with a slit size of 600 μm (77216, Newport) were used for all experiments. The output of the monochromator was reflected downward using an off-axis replicated parabolic mirror with an aluminum coating (50329AL, Newport) to permit the exposure of samples from above.

### UVC characterization and dosimetry

The monochromator spectral output was characterized using a BTS-2048UV Spectroradiometer (Gigahertz-Optik, Inc., Amesbury, MA). With a 600 μm slit width and the 1201.6 g/mm grating, the resolution of the monochromator was 1.9 nm. The measured full width at half maximum was between 2.0 nm and 2.2 nm for all peak wavelengths used in this study. The monochromatic spectral output for wavelengths between 215 nm and 255 nm is shown in Fig. 2 with both a log (panel A) and linear y-axis (panel B). The throughput of the system was measured using an 843-R optical power meter (Newport) with a recently calibrated 818-UV/DB silicon detector (Newport). The total optical power output was measured for each wavelength examined in this work, and this data is plotted in Fig. 3. The irradiance at the target surface was determined by dividing the optical power by the beam area at the exposure plane. The beam area was characterized by using a piece of ultraviolet sensitive film (OrthoChromic Film OC-1, Orthochrome Inc., Hillsborough, NJ) (32). The film was placed at the exposure plane and irradiated to cause a color change illustrating the total exposure area. This area was approximately an 8 mm x 10 mm ellipse, with an area of 62.8 mm^2^. The irradiance for each peak wavelength is also plotted on Fig. 3. The total exposure time for a given wavelength was determined by dividing the desired radiant exposure dose by the irradiance.

**Figure 2.**
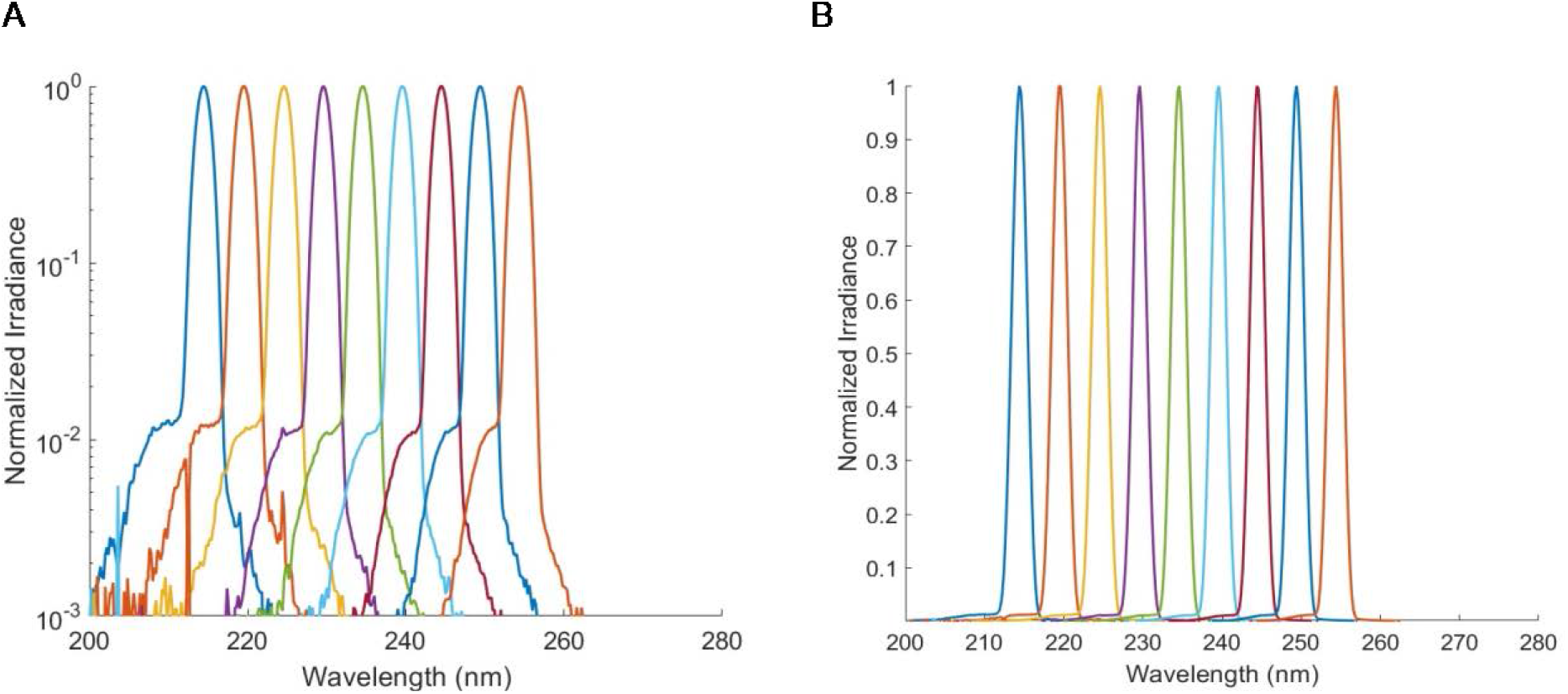
Spectral output of the monochromator for wavelengths tested plotted on a A) log and B) linear scale. The FWHM for each peak wavelength was between 2.0 nm and 2.2 nm.

**Figure 3.**
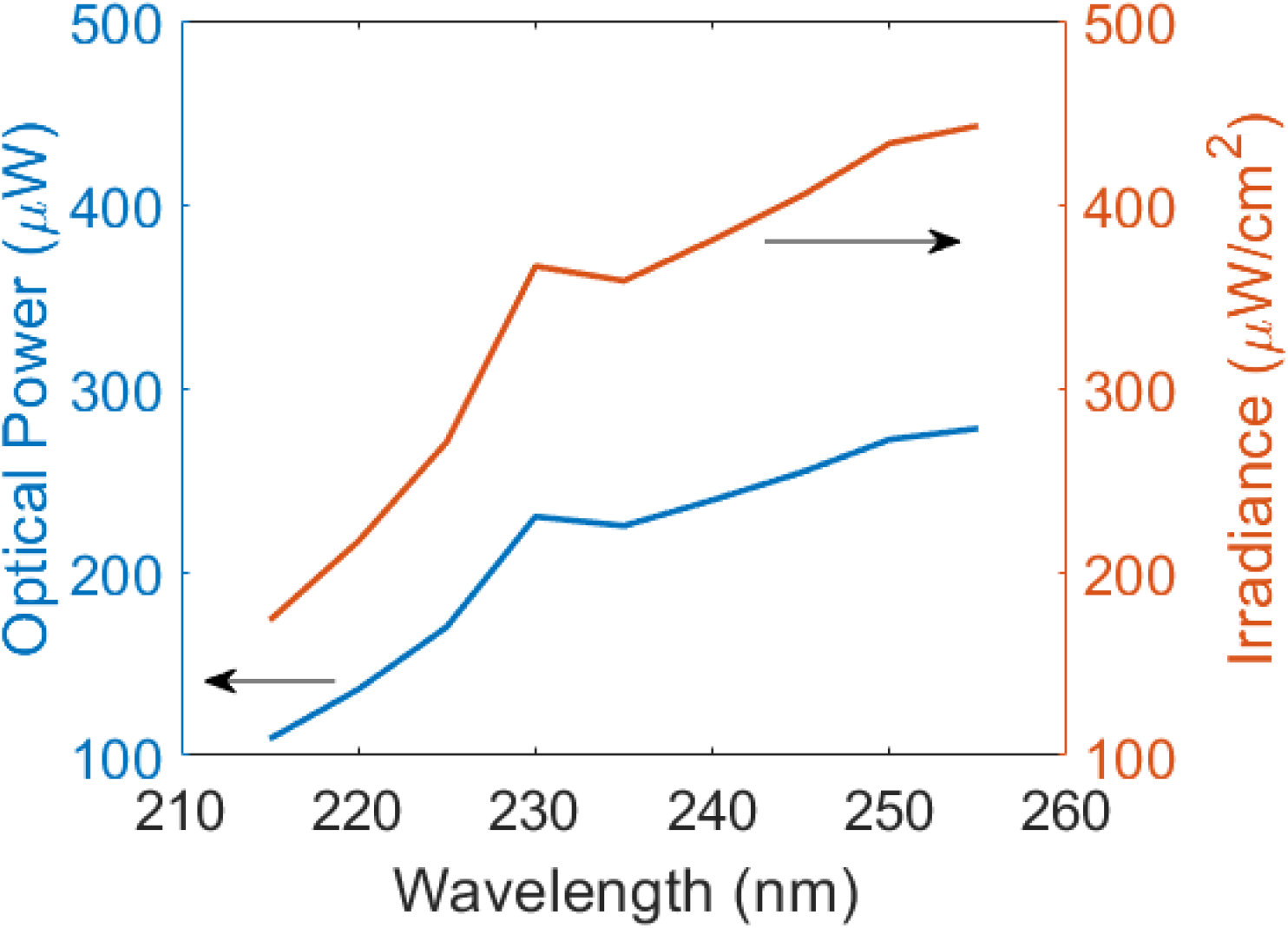
Monochromator optical throughput and irradiance. The total optical power was distributed over an ellipse with an area of 62.8 mm^2^.

### Measurement of UVC-induced CPD epidermal lesions in a 3-D human skin model

We used the 3-D human skin model EpiDerm-FT (MatTek Corp., Ashland, MA) which is derived from single adult donors. EpiDerm-FT is a full-skin thickness construct that recapitulates the key components of human skin, consisting of 8-12 cell layers of normal human epidermal keratinocytes and dermal fibroblasts that form basal, spinous, granular, and cornified layers analogous to those found *in vivo* (33).

The tissues were exposed to a radiant exposure dose of 100 mJ/cm^2^ using narrow bandwidth exposures centered at wavelength of 215, 220, 225, 230, 235, 240, 245, 250 or 255 nm. Experimental controls were unexposed 3-D tissues. Both the sham (controls) and exposed tissues were fixed 15 min after exposure. Two tissues were exposed at each of the examined wavelengths, and we measured the percentage of the most abundant premutagenic DNA photolesion, cyclobutane pyrimidine dimers (CPD) (34), in epidermal keratinocytes, analyzing multiple fields within each tissue. The CPDs were detected using a standard immunohistochemical method previously described (35).

For each tissue, multiple randomly-selected fields of view were analyzed across the tissues to determine the CPD incidence in the different strata of the epidermis (stratum granulosum, stratum spinosum, and stratum basale, see Fig. 1), as well as averaged over the entire epidermis. CPD yields represent the average ± standard deviation of keratinocytes exhibiting dimers divided by the total number of cells measured in a randomly selected fields of view. A typical field of view is shown in Fig. 1, and the total number of cells were determined by counting the number of nuclei positive for 4’,6-diamidino-2-phenylindole (DAPI) using the coverslip mounting medium with DAPI (Vectashield, Burlingame, CA). Similarly, the percentage of CPD-positive keratinocytes in each layer of the epidermis was obtained by dividing the number of positive cells in that layer by the total number of cells counted in that specific layer. Uncertainties (95% and 99% confidence intervals) for the percentage of CPD positive cells were estimated for each sample based on Agresti-Coull (adjusted Wald) confidence interval analysis (36).

## RESULTS AND DISCUSSION

We irradiated the 3-D skin model with narrow bandwidth UVC exposures, in order to examine changes in DNA damage biological effects associated with small changes in wavelength. With a full width half maximum between 2.0 nm and 2.2 nm for all peak wavelengths used in this study, we exposed multiple 3-D models of normal human skin to 100 mJ/cm^2^ of narrow bandwidth UVC at nine different wavelengths from 215 nm to 255 nm (215, 220, 225, 230, 235, 240, 245, 250, 255 nm). The exposure of 100 mJ/cm^2^ was chosen to be somewhat larger than the current Threshold Limit Value / Exposure Limit for 222 nm of 23 mJ/cm^2^ for an 8-hour exposure.

After irradiation, sample preparation and staining we analyzed multiple fields throughout the epidermis for CPD lesions, at the superficial granular layer (stratum granulosum), at intermediate depths (stratum spinosum) and at the basal cell layer (stratum basale).

At the five far-UVC wavelengths that we studied (215, 220, 225, 230, 235 nm), we analyzed a total of 76 fields throughout the epidermis, with an average of 95 keratinocyte cells per field. The results are summarized in Fig. 4A. Based on Agresti-Coull (adjusted Wald) confidence interval analysis (36), in none of the 76 epidermal fields in the far-UVC exposed samples did we observe a statistically significant increase in CPD photolesions relative to zero exposure controls.

**Figure 4.**
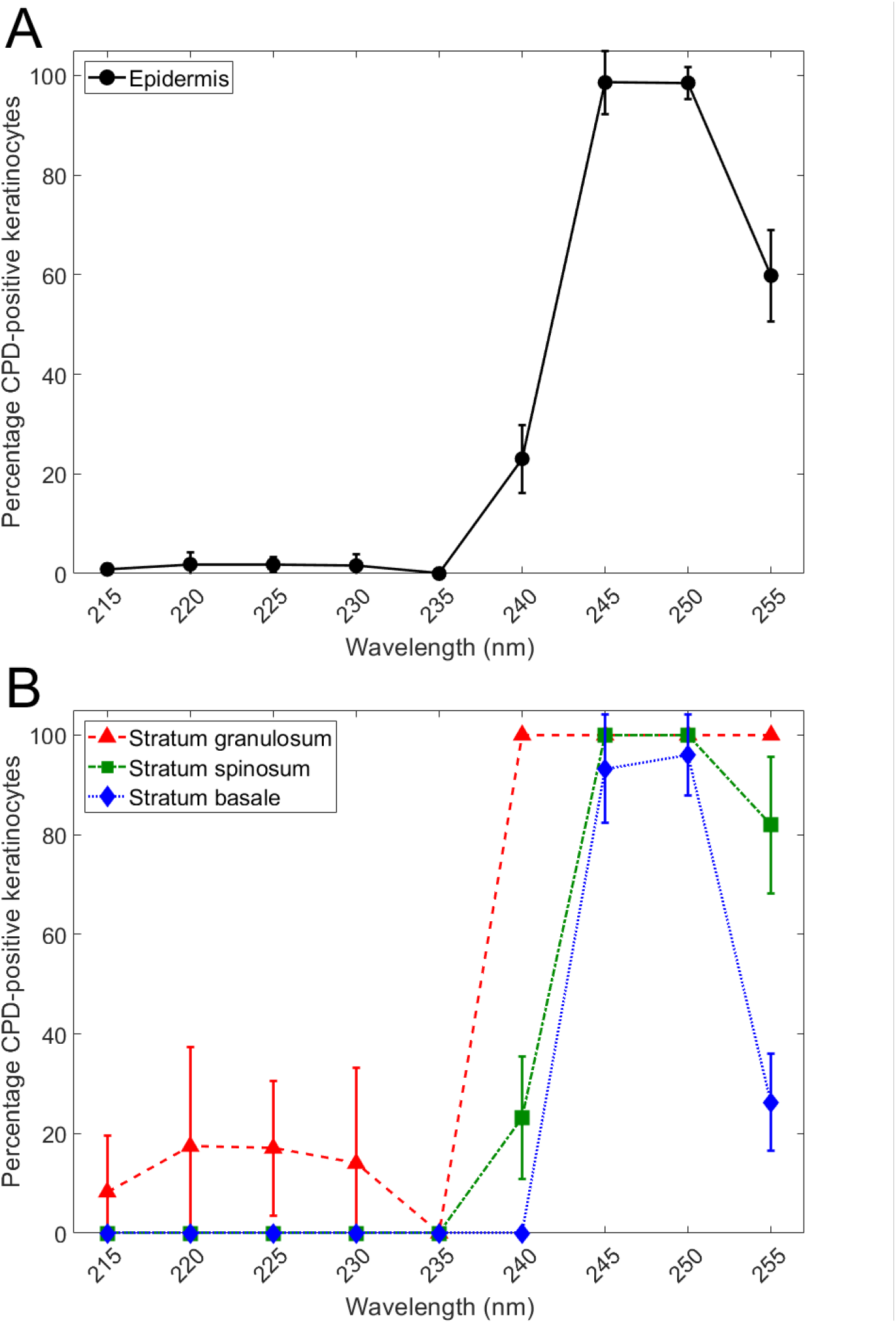
Percentage of DNA photodamage induced by 100 mJ/cm^2^ in the UVC wavelength range. Percentage of the total keratinocytes positive for CPD counted in A) the whole epidermis and B) each layer (see Fig. 1)of the epidermis. Error bars indicate standard deviations.

At the four higher UVC wavelengths that we studied (240, 245, 250, 255 nm), we analyzed a total of 40 fields throughout the epidermis, with an average of 109 keratinocyte cells analyzed per field. The results are summarized in Fig. 4A, and in contrast to the far-UVC results at 215 to 235 nm, in every one of the 40 epidermal fields observed after 240 to 255 nm exposure, a statistically significant increase in CPD photolesions relative to controls was observed, again based on Agresti-Coull confidence interval analysis.

Fig. 4B shows the same CPD data but broken down into the three epidermal strata (see Fig. 1), the stratum granulosum, the stratum spinosum and the stratum basale. As shown in Fig. 4B, in the far-UVC wavelength range (215 to 235 nm) no CPD lesions were observed in either the stratum spinosum or the stratum basale, but a significant increase in CPDs was observed in the superficial stratum granulosum. By contrast, at the higher UVC wavelengths (240 to 255 nm), significant increases in CPDs vs. controls were observed in all layers, except in the basal layer at 240 nm.

To put these stratum-specific results into context (and see Fig. 1), the stratum basale is the deepest layer of the epidermis, where basal cells, including melanocytes, are constantly dividing and migrating upwards; above the stratum basale is the stratum spinosum which contains squamous cells and provides the skin’s structural integrity; and above the stratum spinosum is the stratum granulosum which contains dead or dying cells whose nuclei and other organelles are disintegrating as the cells move up into the stratum corneum (37). Thus from a long-term safety perspective, the concern relates to DNA damage to cells in the stratum basale and stratum spinosum, which contain living basal cells, melanocytes and squamous cells (24, 25, 38). DNA damage to cells in the stratum granulosum or, of course, the stratum corneum is of much less concern, as these contain dead or dying cells.

We may conclude from these results that, at UVC exposures of 100 mJ/cm^2^, far-UVC (215 to 235 nm) did not produce a significant increase in photodamage averaged over the epithelium, and did not produce any photodamage in the relevant epithelial layers, namely the stratum basale and the stratum spinosum. By contrast, exposure to the higher UVC wavelengths studied (240 to 255 nm) does produce significant increases in photodamage in the epithelium, and at each of the epithelial layers studied.

As well as providing support for the basic concept of far-UVC safety, the results shown here should allow for optimized design of UVC filters designed to reduce the higher-wavelength UVC spectral impurities that are typically associated with far-UVC light sources (17). In addition, these results should contribute towards improved recommendations of UVC action spectra, currently under review by ACGIH (4); these results suggest that, at least for skin, the currently recommended Threshold Limit Values for far-UVC may be overprotective.

In conclusion, these results provide quantitative wavelength-specific data supporting the safe use of far-UVC in occupied public settings. The data were generated using a realistic 3-D human skin model exposed to UVC exposures of 100 mJ/cm^2^, somewhat higher than the current Threshold Limit Value / Exposure Limit of 23 mJ/cm^2^ / 8 hour exposure. At this exposure no photodamage was observed in the key epidermal layers of the stratum basale and the stratum spinosum – the locations of epidermal basal cells, melanocytes and squamous cells - at the far-UVC wavelengths of 215, 220, 225, 230 and 235 nm, in contrast to higher UVC wavelengths (240, 245, 250 and 255 nm) where significant levels of photodamage were observed.

## Acknowledgments

This work funded in part under an AFWERX SBIR with the 189th Airlift Wing, Arkansas Air National Guard and Far UV Technologies, as well as from the Shostack Foundation and LumenLabs. We thank Dr. Gerhard Randers-Pehrson for his conceptual insights. We are grateful to the Ultraviolet Radiation Group of the Sensor Science Division in the NIST Physical Measurement Laboratory for assistance with the monochromator system design.

## Data sharing

Data are available on the Open Science Framework (OSF) repository: https://osf.io/aj7yc/

## REFERENCES

1. Kowalski, W.J., Ultraviolet Germicidal Irradiation Handbook: UVGI for Air and Surface Disinfection. 2009: New York: Springer.

2. Wells, W., M. Wells, and T. Wilder, The environmental control of epidemic contagion. I. An epidemiologic study of radiant disinfection of air in day schools. Am J Hyg, 1942. 35: p. 97–121.

3. Nardell, E., R. Vincent, and D.H. Sliney, Upper-room ultraviolet germicidal irradiation (UVGI) for air disinfection: a symposium in print. Photochem Photobiol, 2013. 89(4): p. 764–9.

4. 2021 Threshold Limit Values and Biological Exposure Indices. 2021: American Conference of Governmental Industrial Hygienists.

5. The International Commission on Non-Ionizing Radiation Protection, Guidelines on limits of exposure to ultraviolet radiation of wavelengths between 180 nm and 400 nm (incoherent optical radiation). Health Physics, 2004. 87(2): p. 171–186.

6. Buchan, A.G., L. Yang, and K.D. Atkinson, Predicting airborne coronavirus inactivation by far-UVC in populated rooms using a high-fidelity coupled radiation-CFD model. Scientific Reports, 2020. 10(1): p. 19659.

7. Buonanno, M., B. Ponnaiya, D. Welch, M. Stanislauskas, G. Randers-Pehrson, L. Smilenov, F.D. Lowy, D.M. Owens, and D.J. Brenner, Germicidal Efficacy and Mammalian Skin Safety of 222-nm UV Light. Radiation Research, 2017. 187(4): p. 483–491.

8. Buonanno, M., D. Welch, I. Shuryak, and D.J. Brenner, Far-UVC light (222 nm) efficiently and safely inactivates airborne human coronaviruses. Scientific Reports, 2020. 10(1): p. 10285.

9. Eadie, E., W. Hiwar, L. Fletcher, E. Tidswell, P. O’Mahoney, M. Buonanno, D. Welch, C.S. Adamson, D.J. Brenner, C. Noakes, and K. Wood, Far-UVC efficiently inactivates an airborne pathogen in a room-sized chamber. Scientific Reports www.researchsquare.com/article/rs-908156/v1. Submitted.

10. Glaab, J., N. Lobo-Ploch, H.K. Cho, T. Filler, H. Gundlach, M. Guttmann, S. Hagedorn, S.B. Lohan, F. Mehnke, and J. Schleusener, Skin tolerant inactivation of multiresistant pathogens using far-UVC LEDs. Scientific reports, 2021. 11(1): p. 1–11.

11. Goh, J.C., D. Fisher, E.C.H. Hing, L. Hanjing, Y.Y. Lin, J. Lim, O.W. Chen, and L.T. Chye, Disinfection capabilities of a 222 nm wavelength ultraviolet lighting device: a pilot study. Journal of Wound Care, 2021. 30(2): p. 96–104.

12. Kitagawa, H., Y. Kaiki, K. Tadera, T. Nomura, K. Omori, N. Shigemoto, S. Takahashi, and H. Ohge, Pilot study on the decontamination efficacy of an installed 222-nm ultraviolet disinfection device (Care222™), with a motion sensor, in a shared bathroom. Photodiagnosis and Photodynamic Therapy, 2021. 34: p. 102334.

13. Kitagawa, H., T. Nomura, T. Nazmul, R. Kawano, K. Omori, N. Shigemoto, T. Sakaguchi, and H. Ohge, Effect of intermittent irradiation and fluence-response of 222 nm ultraviolet light on SARS-CoV-2 contamination. Photodiagnosis and Photodynamic Therapy, 2021. 33: p. 102184.

14. Kitagawa, H., T. Nomura, T. Nazmul, K. Omori, N. Shigemoto, T. Sakaguchi, and H. Ohge, Effectiveness of 222-nm ultraviolet light on disinfecting SARS-CoV-2 surface contamination. American journal of infection control, 2021. 49(3): p. 299–301.

15. Welch, D., M. Buonanno, V. Grilj, I. Shuryak, C. Crickmore, A.W. Bigelow, G. Randers-Pehrson, G.W. Johnson, and D.J. Brenner, Far-UVC light: A new tool to control the spread of airborne-mediated microbial diseases. Scientific Reports, 2018. 8(1): p. 2752.

16. Barnard, I.R.M., E. Eadie, and K. Wood, Further evidence that far-UVC for disinfection is unlikely to cause erythema or pre-mutagenic DNA lesions in skin. Photodermatol Photoimmunol Photomed, 2020. 36(6): p. 476–477.

17. Buonanno, M., D. Welch, and D.J. Brenner, Exposure of Human Skin Models to KrCl Excimer Lamps: The Impact of Optical Filtering†. Photochemistry and Photobiology, 2021. 97(3): p. 517–523.

18. Eadie, E., I.M. Barnard, S.H. Ibbotson, and K. Wood, Extreme exposure to filtered far-UVC: A case study. Photochemistry and Photobiology, 2021. 97(3): p. 527–531.

19. Fukui, T., T. Niikura, T. Oda, Y. Kumabe, H. Ohashi, M. Sasaki, T. Igarashi, M. Kunisada, N. Yamano, K. Oe, T. Matsumoto, T. Matsushita, S. Hayashi, C. Nishigori, and R. Kuroda, Exploratory clinical trial on the safety and bactericidal effect of 222-nm ultraviolet C irradiation in healthy humans. PLOS ONE, 2020. 15(8): p. e0235948.

20. Hickerson, R., M. Conneely, S. Hirata Tsutsumi, K. Wood, D. Jackson, S. Ibbotson, and E. Eadie, Minimal, superficial DNA damage in human skin from filtered far-ultraviolet C. British Journal of Dermatology, 2021.

21. Finlayson, L., I.R.M. Barnard, L. McMillan, S.H. Ibbotson, C.T.A. Brown, E. Eadie, and K. Wood, Depth Penetration of Light into Skin as a Function of Wavelength from 200 to 1000 nm. Photochemistry and Photobiology, 2021.

22. Cadet, J., Harmless Effects of Sterilizing 222-nm far-UV Radiation on Mouse Skin and Eye Tissues. Photochemistry and Photobiology, 2020. 96(4): p. 949–950.

23. Cesarini, J.-P., C. Cole, and F. de Gruijl, UV-C photocarcinogenesis risks from germicidal lamps. Int Commission Illumination, 2010. 187: p. 1–14.

24. Forbes, P.D., C.A. Cole, and F. de Gruijl, Origins and Evolution of Photocarcinogenesis Action Spectra, Including Germicidal UVC(dagger). Photochem Photobiol, 2021. 97(3): p. 477–484.

25. de Gruijl, F.R. and C.P. Tensen, Pathogenesis of skin carcinomas and a stem cell as focal origin. Frontiers in medicine, 2018. 5: p. 165.

26. Ma, B., P.M. Gundy, C.P. Gerba, M.D. Sobsey, and K.G. Linden, UV Inactivation of SARS-CoV-2 across the UVC Spectrum: KrCl* Excimer, Mercury-Vapor, and Light-Emitting-Diode (LED) Sources. Appl Environ Microbiol, 2021. 87(22): p. e0153221.

27. Eadie, E., P. O’Mahoney, L. Finlayson, I.R.M. Barnard, S.H. Ibbotson, and K. Wood, Computer Modeling Indicates Dramatically Less DNA Damage from Far-UVC Krypton Chloride Lamps (222 nm) than from Sunlight Exposure. Photochemistry and Photobiology, 2021.

28. Hanamura, N., H. Ohashi, Y. Morimoto, T. Igarashi, and Y. Tabata, Viability evaluation of layered cell sheets after ultraviolet light irradiation of 222 nm. Regenerative Therapy, 2020. 14: p. 344–351.

29. Kaidzu, S., K. Sugihara, M. Sasaki, A. Nishiaki, T. Igarashi, and M. Tanito, Evaluation of acute corneal damage induced by 222-nm and 254-nm ultraviolet light in Sprague–Dawley rats. Free Radical Research, 2019. 53(6): p. 611–617.

30. Kaidzu, S., K. Sugihara, M. Sasaki, A. Nishiaki, H. Ohashi, T. Igarashi, and M. Tanito, Re-Evaluation of Rat Corneal Damage by Short-Wavelength UV Revealed Extremely Less Hazardous Property of Far-UV-C. Photochemistry and Photobiology, 2021. 97(3): p. 505.

31. Yamano, N., M. Kunisada, S. Kaidzu, K. Sugihara, A. Nishiaki-Sawada, H. Ohashi, A. Yoshioka, T. Igarashi, A. Ohira, M. Tanito, and C. Nishigori, Long-term Effects of 222-nm ultraviolet radiation C Sterilizing Lamps on Mice Susceptible to Ultraviolet Radiation. Photochem Photobiol, 2020. 96(4): p. 853–862.

32. Welch, D. and D.J. Brenner, Improved Ultraviolet Radiation Film Dosimetry Using OrthoChromic OC-1 Film(dagger). Photochemistry and Photobiology, 2021. 97(3): p. 498–504.

33. Kubilus, J., P.J. Hayden, S. Ayehunie, S.D. Lamore, C. Servattalab, K.L. Bellavance, J.E. Sheasgreen, and M. Klausner, Full Thickness EpiDerm: a dermal-epidermal skin model to study epithelial-mesenchymal interactions. Altern Lab Anim, 2004. 32 Suppl 1A: p. 75–82.

34. Pfeifer, G.P. and A. Besaratinia, UV wavelength-dependent DNA damage and human non-melanoma and melanoma skin cancer. Photochem Photobiol Sci, 2012. 11(1): p. 90–7.

35. Buonanno, M., M. Stanislauskas, B. Ponnaiya, A.W. Bigelow, G. Randers-Pehrson, Y. Xu, I. Shuryak, L. Smilenov, D.M. Owens, and D.J. Brenner, 207-nm UV Light-A Promising Tool for Safe Low-Cost Reduction of Surgical Site Infections. II: In-Vivo Safety Studies. PLoS One, 2016. 11(6): p. e0138418.

36. Agresti, A. and B.A. Coull, Approximate is better than “exact” for interval estimation of binomial proportions. The American Statistician, 1998. 52(2): p. 119–126.

37. Barbieri, J.S., K. Wanat, and J. Seykora, Skin: Basic Structure and Function, in Pathobiology of Human Disease, L.M. McManus and R.N. Mitchell, Editors. 2014, Academic Press: San Diego. p. 1134–1144.

38. Black, H.S., F.R. deGruijl, P.D. Forbes, J.E. Cleaver, H.N. Ananthaswamy, E.C. deFabo, S.E. Ullrich, and R.M. Tyrrell, Photocarcinogenesis: an overview. J Photochem Photobiol B, 1997. 40(1): p. 29–47.

